# Secreted and cell surface proteome analysis of pathogenic *Corynebacterium diphtheriae* reveals proteins relevant to virulence

**DOI:** 10.1101/2020.09.06.285155

**Authors:** Andrew K. Goring, Yu Chen, Robert T. Clubb, Joseph A. Loo

## Abstract

*Corynebacterium diphtheriae* is responsible for the severe upper respiratory tract disease, diphtheria, which can be fatal in healthy individuals without proper treatment. Its interaction with the infected host is crucial to its virulence, secreting diphtheria toxin and a variety of machinery to acquire nutrients for further import into the cell. Additionally, it shares a conserved iron import mechanism with related pathogenic actinobacteria such as *Mycobacterium tuberculosis*, and contains sortase-mediated anchored cell wall proteins similar to other gram-positive bacteria. Information obtained from secreted and cell-surface proteomes are relevant for the study of diphtheria and related bacterial infections. Mass spectrometry-based proteomics measurements on samples of pathogenic *Corynebacterium diphtheriae* culture supernatant and cell-surface digested proteins identified greater than 3 times more proteins than a previous similar study. The diphtheria toxin was identified, as well as pathologically relevant proteins involved in iron-uptake and cell adhesion. For instance, 17 proteins predicted to be on the iron-regulated DtxR promoter were identified in cells grown under different iron concentrations, opening the door to performing comparative quantitative proteomics studies on samples where iron sources are modulated. Results of this study serve as a repository for future studies on pathogenic *C. diphtheriae* uptake.

## Introduction

*Corynebacterium diphtheriae* (*C. diphtheriae*) is responsible for the severe upper-respiratory tract disease, diphtheria. The diphtheria toxin (DT) produced during an infection causes systemic effects such as myocarditis and neuropathy, associated with increased risk of fatality.^1^ *C. diphtheriae* belongs to the phylum actinobacteria, which contains a large group of medically and industrially important Gram-positive bacteria including *Mycobacterium tuberculosis, Nocardia abscessus, Propionibacterium* spp., *Tropheryma* spp., and *Bifidobacterium* spp. More closely related species such as *Corynebacterium ulcerans* and *Corynebacterium pseudotuberculosis* species can also produce DT. In addition to toxigenic, DT-producing *C. diphtheriae*, there are non-toxigenic strains that do not produce the toxin but can still mount fatal infections in healthy individuals.

In order to proliferate, *C. diphtheriae* requires iron. It is an essential metal cofactor used to perform a wide variety of important cellular processes such as DNA replication, electron transport, and amino acid synthesis. Free iron is limiting in the host, with 75-80% of the body’s iron bound by hemoglobin (Hb) in the form of heme (iron–protoporphyrin IX).^2,3^ Thus, *C. diphtheriae* has developed elaborate systems to acquire iron from a host environment. Genes encoding components of the heme-uptake system are expressed from five genetic loci in iron-deplete conditions (*hmu, chtA-chtB, cirA-chtC, hbpA* and *hmuO*).^4^ Iron levels also control the amount of DT that is produced. DT is secreted and eventually enters nearby host cells and inhibits protein synthesis, ultimately leading to cell death.^5^ *C. diphtheriae*’s virulence and production of DT were shown to be controlled by iron levels and its promoter is therefore named diphtheria toxin repressor (DtxR).^6^ Protein machinery associated with iron-uptake, such as the heme-import machinery, is also controlled under the DtxR regulon. Vaccines aimed at *C. diphtheriae* have primarily targeted DT, but other proteins conserved across strains found in *C. diphtheriae* vaccines could also reside on the surface and be immunogenic.^7^ Virulence has also been shown to be associated with cell surface anchoring proteins in *C. diphtheriae* and others in Gram-positive bacteria.^8^ Thus, knowledge of the cell surface proteome of *C. diphtheriae* is important for vaccine development and identifying new drug targets.

From a 2006 proteomics study^9^ of *C. diphtheriae* using 2D gel electrophoresis separation of *C. diphtheriae* proteins, 85 surface and secreted proteins were identified. However, more recent studies on *Corynebacterium ulcerans* and *Corynebacterium glutamicum* identified many more proteins on or predicted to be localized to the cell surface,^10–12^ suggesting that more coverage of the secreted and cell surface *C. diphtheriae* proteome should be obtainable. We used a high resolution HPLC mass spectrometry experimental strategy (e.g., trypsin digestion of precipitated supernatant and intact cells) to obtain an updated secretome and surfaceome of *C. diphtheriae*.

The strain studied in the present study is wild-type *C. diphtheriae* strain 1737, a clinical isolate from the early 1990’s outbreak in Russia,^13^ which encodes for the *tox* gene and is relevant to clinical research. We compare the results of this bacteria grown under minimal iron conditions to that of the 2006 study, and search for proteins related to virulence such as DtxR-regulated proteins and surface-anchored proteins. We also grew strains in high iron conditions and compared identified DtxR-regulated proteins, with a focus on identifying known heme uptake components, to gain preliminary insight on how the bacteria may change its extracellular expression levels under stress conditions.

## Methods

### Cell Culture

Strains were streaked onto heart-infusion broth agar plates and grown at 37 °C until colonies were formed. Individual colonies were used to inoculate 10 mL cultures in minimal media (mPGT) with 1.0 µM FeCl_3_ supplemented for growth overnight. Sample growth was measured spectroscopically by measuring their optical density at 600 nm (OD_600_). Cultures were then spun down, and washed with iron-free mPGT media. The samples were resuspended and used to inoculate 10 mL cultures in minimal mPGT media with 0.3 µM (low iron) or 10 µM (high iron) FeCl_3_ supplemented with a starting OD_600_ of 0.01. The samples were harvested at mid-log phase (OD_600_ between 1 and 2).

### Isolation of Secreted Proteins

Cultures were spun down and the supernatant separated, spun down again, and filtered to ensure it was cell-free. The extracellular proteins in the filtrate were then concentrated in a 3k MWCO Amicon centrifugal filter, disulfide-reduced, Cys-alkylated, and precipitated with acetone. Precipitated proteins were then pelleted, washed with acetone, dried and prepared for MS analysis by trypsin digestion and de-salting with a C18-packed StageTip.^14^

### Isolation of Surface Proteins

Cultures were spun down and the pellet was used for cell surface preparation after removing the supernatant. It was washed 3 times with PBS buffer, pH 7.4, before being digested directly on the intact cell wall with trypsin overnight. Disulfide reduction and Cys-alkylation were performed on the resulting peptides after cells were removed by centrifugation and filtration.

### LC MS/MS

A sample volume of 1.0 µL was injected to an Ultimate 3000 nano LC, which was equipped with a 75 µm x 2 cm trap column packed with C18 3µm bulk resins (Acclaim PepMap 100, Thermo Scientific) and a 75 µm x 15 cm analytical column with C18 2 µm resins (Acclaim PepMap RSLC, Thermo Scientific). The nanoLC gradient was 3-35% solvent B (A = H_2_O with 0.1% formic acid; B = acetonitrile with 0.1% formic acid) over 40 min and from 35% to 85% solvent B in 5 min at flow rate 300 nL/min. The nanoLC was coupled to a Q Exactive Plus orbitrap mass spectrometer (Thermo Fisher Scientific, San Jose, CA). The ESI voltage was set at 1.9 kV, and the capillary temperature was set at 275 ^°C^. Full spectra (*m/z* 350 - 2000) were acquired in profile mode with resolution 70,000 at *m/z* 200 with an automatic gain control (AGC) target of 3 × 10^6^. The most abundant 15 ions per scan were subjected to fragmentation by higher-energy collisional dissociation (HCD) with normalized collisional energy of 25. MS/MS spectra were acquired in centroid mode with resolution 17,500 at *m/z* 200. The AGC target for fragment ions was set at 2 × 10^4^ with a maximum injection time of 50 ms. Charge states 1, 7, 8, and unassigned were excluded from tandem MS experiments. Dynamic exclusion was set at 45.0 s.

### Data Analysis

Raw MS data was searched against the Uniprot human database by Proteome Discovered version 1.4. The following parameters were set: precursor mass tolerance ±10 ppm, fragment mass tolerance ±0.02 Da for HCD, up to two miscleavages by trypsin, methionine oxidation as variable modification. The false discovery rate was at 1.0% and a minimum of 2 peptides was required for protein identification.

## Results and Discussion

### Protein Identifications

We set out to compare the surface and secreted proteomes of *C. diphtheriae* under iron-limiting conditions (0.3 µM) to that of previous proteomics studies, to assess consistency with literature and improvements upon number of proteins identified. A total of 310 unique proteins from *C. diphtheriae* grown to mid-log phase in minimal media with 0.3 µM supplemented iron were identified from the secreted and cell surface fractions, which had 219 and 229 protein identifications, respectively. There was an overlap of 138 out of 310 total proteins (44%) found in both the secreted and cell surface fractions. This is within the range of the surface/secreted overlap in the 2006 study (29%) and in the *C. ulcerans* study (41% for strain 809 and 58% for strain BR-AD22). Reasons for overlap in the fractions could be due to proteins residing in both fractions, and/or secretion of proteins during the cell surface trypsin digest preparation.

There was an increase in proteins identified and good agreement with the previous 2006 study by Hansmeier et al.^9^ Overall, 247 more proteins were identified in this study than in the 2006 work and 79% of the 2006 protein identifications overlapped with those found in this analysis (**Figure 1**). Of the 35 cell surface proteins identified in the 2006 study, 28 (80%) were identified in this data, and of the 74 secreted proteins found in the 2006 study, 69 (80%) were identified in this data. Interestingly, there were 18 proteins identified in the 2006 study that were not identified in ours. This could be explained by the use of different strains and different growth conditions (LB broth vs. minimal media with supplemented iron; late-log phase sampling vs. mid-log phase sampling).

**Figure 1:**
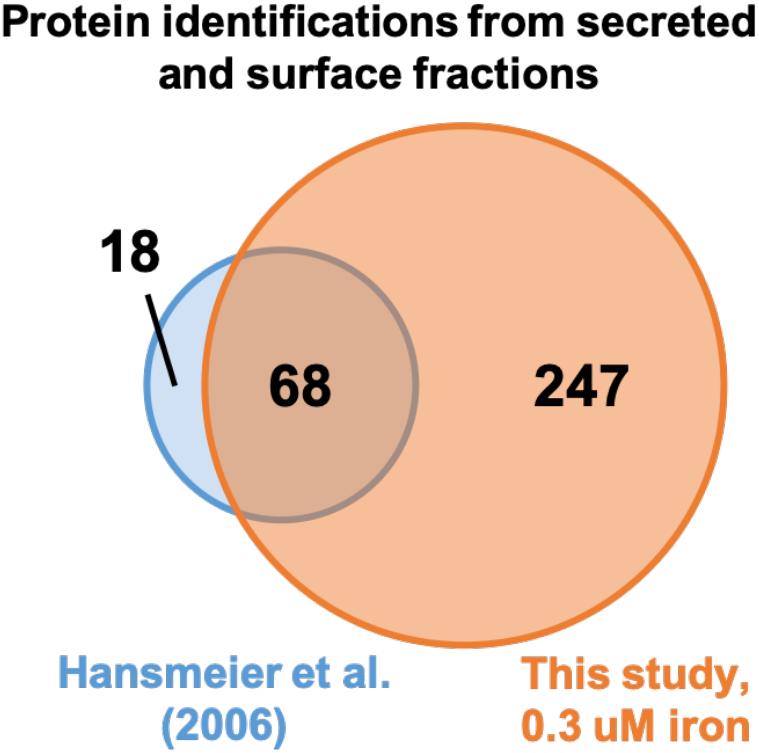
Venn diagram comparing secreted and cell surface fraction protein identifications from *C. diphtheriae* from the 2006 analysis by Hansmeier et al.^9^ and this study (2020).

### Identification of Toxin and Proteins Related to C. diphtheriae Virulence

There are multiple proteins that are implicated in the virulence of *C. diphtheriae* that we looked for in order to evaluate our data set’s significance. First, there is DT, encoded by the *tox* gene (DIP0222) and directly contributing to toxicity. This was identified in the secreted fraction in our study. It was not identified in the previous study as they used a DT knockout strain, *C. diphtheriae* C7_s_(-)^*tox-*^.

The DtxR regulon controls expression of DT and the iron import machinery such as the heme uptake system components, which are required for *C. diphtheriae* to steal hemoglobin from the host. A total of 17 proteins (**Table 1**) were found that are presumed to be under DtxR control from a 2004 genomic analysis by Yellaboina et al.^15^ The previous 2006 study identified 5 proteins under DtxR control (**Table 1**), which were also observed in our analysis. Interestingly, the machinery known to be responsible for acquiring heme and secreted from the host cells were identified in this study (ChtA, ChtB, ChtC, HtaA, HtaB, HbpA, and HmuT), showing the depth of pathway identification in our data set.

**Table 1:** Protein identifications predicted to be controlled by the DtxR regulon by a genomic analysis from Yellaboina et al. (2004).^15^ The export mechanism predicted by SignalP and SecP algorithms is shown in the far right-hand column.

| Gene name | Accession (NCTC 13129) | Study identified* | Description | Export mechanism** |
| --- | --- | --- | --- | --- |
| DIP0543 | Q6NJ69 | 2020 and 2006 | Putative sialidase | SP(Sec/SPI) |
| DIP0544 | Q6NJ68 | 2020 and 2006 | Putative secreted protein | SP(Sec/SPI) |
| *adk*<br>DIP0541 | Q6NJ71 | 2020 and 2006 | Adenylate kinase | N/A |
| *hmuT*<br>DIP0626 | Q6NIZ0 | 2020 and 2006 | Iron-related transport system receptor protein | LIPO(Sec/SPII) |
| *infA*<br>DIP0545 | P61685 | 2020 and 2006 | Translation initiation factor IF-1 | N/A |
| DIP2162 | Q6NEV3 | 2020 high and low iron | Putative ABC transport system, solute-binding protein | TAT(Tat/SPI) |
| DIP2272 | Q6NEK1 | 2020 high and low iron | Possible sortase-like protein | Unknown |
| *rpsM*<br>DIP0546 | Q6NJ66 | 2020 high and low iron | 30S ribosomal protein S13 | Unknown |
| *rplQ*<br>DIP0550 | Q6NJ62 | 2020 high and low iron | 50S ribosomal protein L17 | N/A |
| *htaA*<br>DIP0625 | Q6NIZ1 | 2020 high and low iron | Putative membrane protein | SP(Sec/SPI) |
| *chtB*<br>DIP1519 | Q6NGJ4 | 2020 high and low iron | Putative membrane protein | SP(Sec/SPI) |
| *chtA*<br>DIP1520 | Q6NGJ3 | 2020 high and low iron | Putative membrane protein | SP(Sec/SPI) |
| *htaB*<br>DIP0629 | Q6NIY7 | 2020 high and low iron | Membrane protein | SP(Sec/SPI) |
| *hbpA*<br>DIP2330 | Q6NEE5 | 2020 high and low iron | Putative membrane protein | SP(Sec/SPI) |
| DIP0169 | Q6NK68 | 2020 high and low iron | Putative secreted protein | LIPO(Sec/SPII) |
| DIP0173 | Q6NK64 | 2020 high and low iron | Putative membrane protein | LIPO(Sec/SPII) |
| DIP2331 | Q6NEE4 | 2020 high and low iron | Putative aldehyde dehydrogenase | N/A |
| *rpoA*<br>DIP0549 | Q6NJ63 | 2020 high and low iron only | DNA-directed RNA polymerase subunit alpha | N/A |

\* "2020" represents data from the current study, and "2006" represents data reported in the 2006 publication.<sup>9</sup> "Low" and "high" iron represents 0.3 μM or 10 μM FeCl<sub>3</sub>, respectively.
\*\* "Unknown" - predicted to be secreted through SecP but not identified with any known mechanisms from SignalP; "N/A" - it was not predicted to be secreted by either.

Other proteins associated with virulence include adhesins and fimbral associated proteins. Three sortase-like proteins: DIP2272, DIP2012, and DIP0236 were identified in the 2006 study. Of these, we identified only two (DIP2272 and DIP2012). Interestingly, the unidentified protein, DIP2036, resides on a pathogenicity island (PAI) region between DIP0223-DIP0244.^16^ PAI’s are regions on the genome prone to horizontal gene transfer and are known to contribute to genetic variation within related isolates of *C. diphtheriae* and harbor proteins responsible for virulence.^16^ It is possible that strain variation caused us not to find this protein. Sortase-mediated cell-wall anchored proteins are recognized by sortase through an LPxTG motif,^17^ and are also associated with pathogenicity.^8^ We identified a total of 15 predicted LPxTG proteins (**Table 2**), while 3 were found in the previous 2006 study.

**Table 2:** Protein identification information from predicted cell-wall anchored proteins identified in this study and a 2006 study by Hansmeier et al.^9^ The far right column is the prediction of localization from the CW pred algorithm.

| Gene name | Accession (NCTC 13129) | Study identified* | Description | CW pred |
| --- | --- | --- | --- | --- |
| DIP2325 | Q6NF35 | 2020 and 2006 | Putative surface-anchored fimbrial associated protein | Sortase A |
| DIP2227 | Q6NJW5 | 2020 and 2006 | Putative surface-anchored membrane protein | Sortase D |
| DIP2226 | Q6NEF0 | 2020 high and low iron | Putative surface-anchored protein | Sortase D |
| DIP2093 | Q6NEP1 | 2020 high and low iron | Probable surface-anchored fimbrial subunit | Sortase A |
| DIP2066 | Q6NEP2 | 2020 high and low iron | Probable surface-anchored fimbrial subunit | Sortase A |
| DIP2062 | Q6NF16 | 2020 high and low iron | Putative Sdr-family related adhesin | Sortase D |
| DIP2013 | Q6NF39 | 2020 high and low iron | Putative surface anchored protein | Sortase D |
| DIP2011 | Q6NF81 | 2020 high and low iron | Putative surface-anchored fimbrial subunit | Sortase A |
| DIP2010 | Q6NF83 | 2020 high and low iron | Putative surface anchored protein | Sortase D |
| DIP1724 | Q6NF84 | 2020 high and low iron | Putative surface-anchored membrane protein | Sortase A |
| DIP0443 | Q6NG10 | 2020 high and low iron | Putative surface-anchored membrane protein | Sortase D |
| DIP0278 | Q6NJF9 | 2020 high and low iron | Putative surface-anchored membrane protein | Sortase A |
| DIP0238 | Q6NK02 | 2020 high and low iron | Putative surface-anchored fimbrial subunit | Sortase A |
| DIP0237 | Q6NK03 | 2020 high and low iron | Putative surface anchored protein | Sortase D |
| DIP0236 | Q6NK05 | 2020 high and low iron | Putative fimbrial subunit | Sortase A |
| DIP0235 | Q6NK04 | 2006 only | Putative fimbrial associated sortase-like protein | Membrane |

\* "2020" represents data from the current study, and "2006" represents data reported in the 2006 publication.<sup>9</sup> "Low" and "high" iron represents 0.3 μM or 10 μM FeCl<sub>3</sub>, respectively.

### DtxR-regulated Machinery in High and Low Iron Samples

Two data sets were generated from samples grown under either high (10 uM) or low (0.3 uM) iron-supplemented conditions and the heme-uptake machinery and other DtxR-regulated protein identifications were examined in both. Protein identifications in the high and low iron samples from both secreted and cell surface fractions had an overlap of 72% (**Figure 2**).

**Figure 2:**
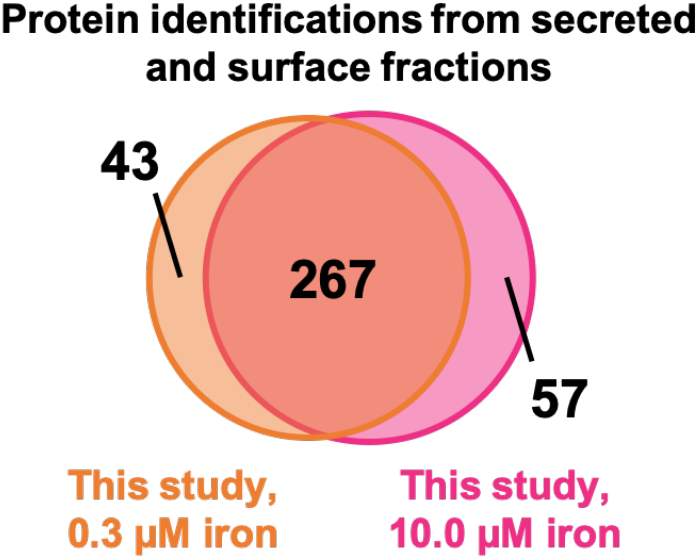
Venn diagram comparing secreted and cell surface fraction protein identifications from *C. diphtheriae* grown under low (0.3 µM) and high (10.0 µM) iron-supplemented conditions.

Furthermore, the differences in protein identifications did not arise significantly from the DtxR-regulated proteins. In both high and low iron data sets, all known heme-transfer components and the same 17 DtxR-regulated proteins were identified (**Table 1**). This opens the doors to performing relative quantification experiments in the future. There was one more DtxR-regulated protein identified in the high-iron condition. It was found to be in the cell surface fraction, which had considerably less fraction of proteins predicted to be “extracellular” (50% compared to 72% in the low iron sample) (**Figure 3**). This protein is not predicted to be extracellular and is annotated as a DNA-directed RNA polymerase subunit alpha, suggesting it could be a product of cell lysis during the cell surface trypsin digest. It would be interesting to see differences in expression levels across these fractions, as was done in whole-cell lysate for *C. glutamicum*.^11^

**Figure 3:**
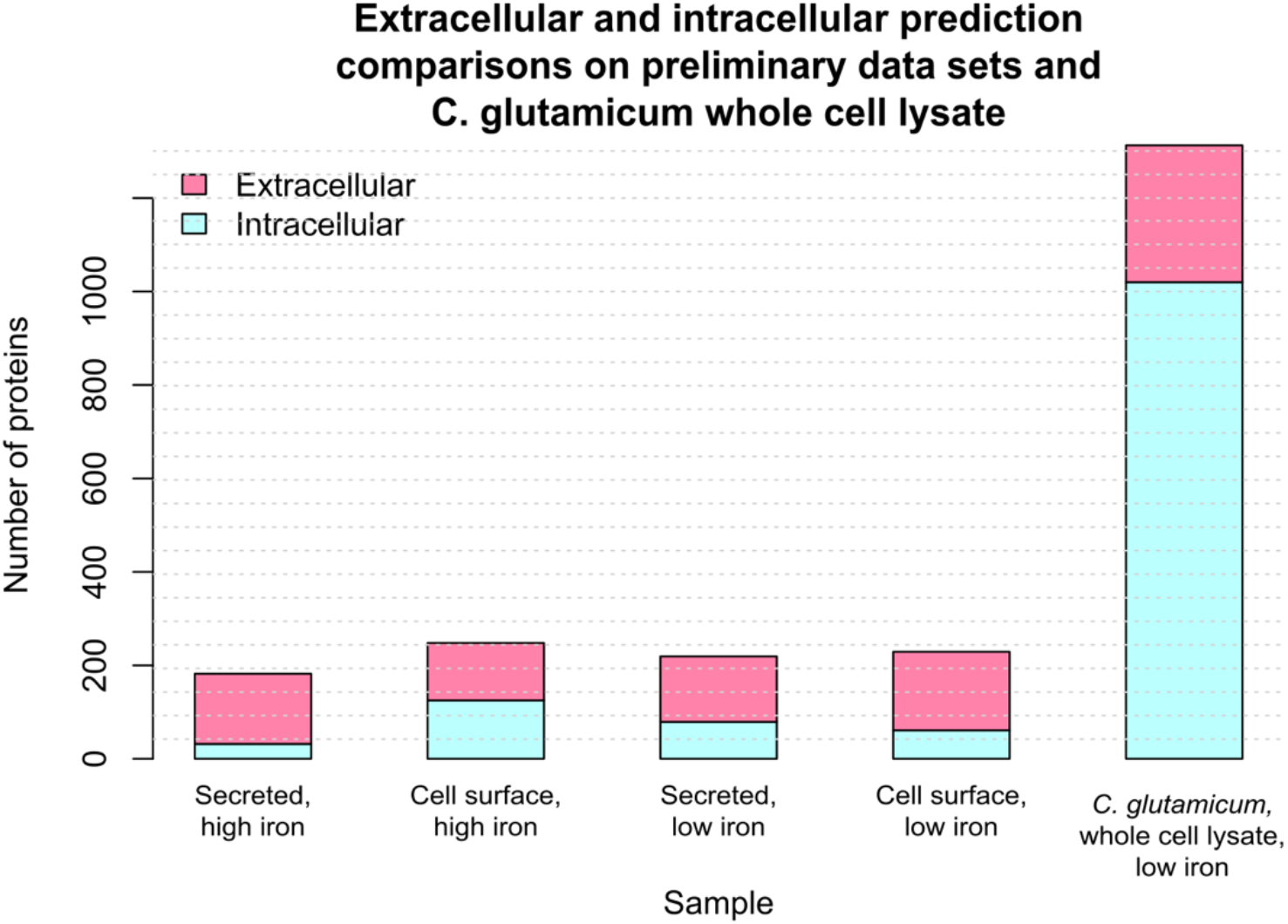
Protein localization analysis on the secreted and cell surface fractions from this study (left four bars) and an example of whole-cell lysate from a closely related homologue, *C. glutamicum*, from Küberl et al.^11^ (right bar). Extracellular predictions are shown in pink and were predicted from SignalP, LipoP, TMHMM, SecretomeP, and CW-pred. The proteins that were not predicted to have extracellular characteristics by any of the algorithms are shown in light blue and categorized as “Intracellular”.

### Cell Surface Localization Comparison

There is selective enrichment of the cell surface proteins based on comparing those identified as extracellular to those from other whole-cell lysate studies, such as that recently done on a related organism, *C. glutamicum*.^11^ An overwhelming majority of the proteins identified in this study were predicted to be membrane associated, cell-wall linked, or secreted (**Figure 3**). A higher number of cell surface predictions for extracellular proteins are in the low iron cell surface sample than in the high iron cell surface sample. This could be indicative of cells being more susceptible to autolysis upon trypsin digestion, perhaps due to physiological differences gained from the different growth conditions. However, more experiments with replicates and experiments with immunoblotting on cytoplasmic control proteins should be performed to confirm this.

### Classes of Protein Identifications

Gene ontology analysis was used to classify the functions of the proteins identified in each fraction from our study. We compared this to the 2006 protein identifications and observed that there were similar trends in categories from cell-surface related proteins, with noticeable magnitude differences to the 2006 study (**Figure 4**). This shows similar protein content was identified, but with more protein identifications to boost the repository of known surface proteins.

**Figure 4:**
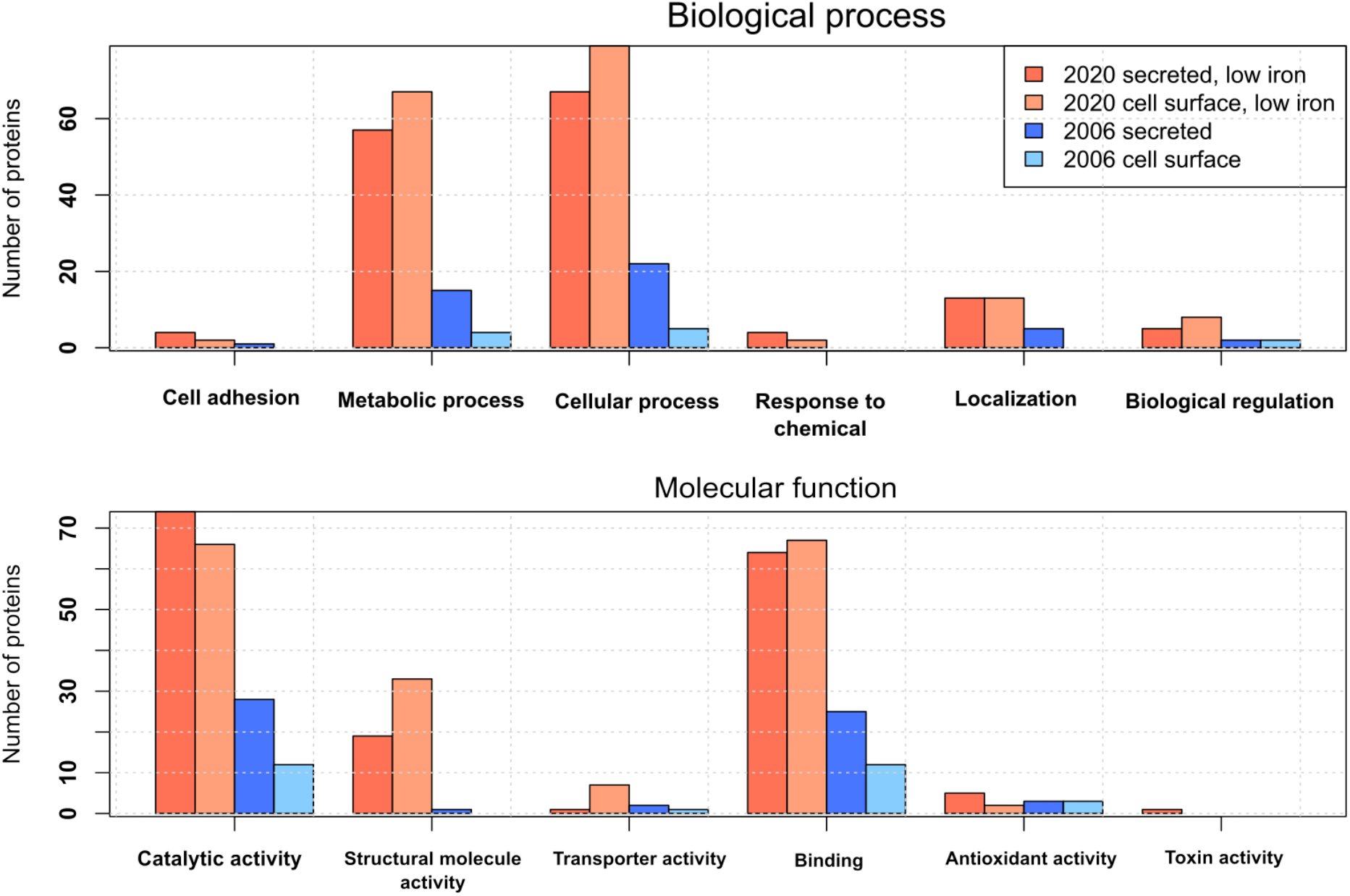
Gene ontology grouping for molecular function and biological process categories from Uniprot. The fractions compared are the cell surface and secreted fractions of the low iron condition from this work (“2020”) and that of the 2006 results by Hansmeier et al.^9^ (“2006”).

### Comment on the Preliminary Data

Only one replicate from each condition was taken. Performing these experiments in biological triplicate could give reliable quantitative information on the heme-uptake machinery and its relative concentrations compared to one-another and across secreted and cell-surface fractions, providing input into its role.

## Conclusion

The cell surface is of high interest because this is the primary means for pathogenic bacteria to interact with its environment. In *C. diphtheriae*, virulence is caused by secretion of DT, and is known to be associated with iron import and pilin assembly for biofilm formation and adherence to host tissue. A data set of cell surface proteins from *C. diphtheriae* was first reported in 2006, identifying 85 proteins from secreted and cell surface samples. In the present study, we obtained a more complete set of genes, 367 in total, from secreted and cell surface proteins. This adds to the previous data set and allows for the filling in of pathways related to virulence and mounting an infection, as observed from looking at DtxR-regulated machinery, LPxTG motif-containing proteins, and a gene ontology analysis. This data can be used for pathway study and vaccine development in the future. Furthermore, there was good agreement in the DtxR-regulated machinery from samples grown at different iron concentrations, where all heme uptake components predicted to be extra-cellular or cell surface associate associated were identified. More detailed relative quantification studies with varying iron concentrations will be performed in the future to understand how *C. diphtheriae* alters expression levels in different fractions to acquire iron from the host during an infection.

## Acknowledgment

This work was supported by the U.S. Department of Energy Office of Science, Office of Biological and Environmental Research program under Award Number DE-FC02-02ER63421 and National Institutes of Health Grants AI52217 (R.T.C.), and GM103479 (J.A.L.). A.K.G. is supported by a Cellular and Molecular Biology Training Grant (Ruth L. Kirschstein National Research Service Award GM007185).

